# PrankWeb: web server for ligand binding-site prediction and visualization

**DOI:** 10.1101/590877

**Authors:** Lukas Jendele, Radoslav Krivak, Petr Skoda, Marian Novotny, David Hoksza

## Abstract

PrankWeb is an online resource providing an interface to P2Rank, a state-of-the-art ligand binding site prediction method. P2Rank is a template-free machine learning method which is based on the prediction of ligandability of local chemical neighborhoods centered on points placed on a solvent accessible surface of a protein. Points with high ligandability score are then clustered to form the resulting ligand binding sites. On top of that, PrankWeb then provides a web interface enabling users to easily carry out the prediction and visually inspect the predicted binding sites via an integrated sequence-structure view. Moreover, PrankWeb can determine sequence conservation for the input molecule and use it in both the prediction and results visualization steps. Alongside its online visualization options, PrankWeb also offers the possibility to export the results as a PyMOL script for offline visualization. The web frontend communicates with the serer side via a REST API. Therefore, in high-throughput scenarios users can utilize the server API directly, bypassing the need for a webbased front end or installation of the P2Rank application. PrankWeb is available at http://prankweb.cz/. The source code of the web application and the P2Rank method can be accessed at https://github.com/jendelel/PrankWebApp and https://github.com/rdk/p2rank, respectively.

## INTRODUCTION

The field of structural biology has recently experienced an enormous progress in all aspects of structural determination process and as a result 3D structures of proteins are becoming increasingly available. Structural genomics consortia are even solving structures of proteins without known functions (1). The information acquired from 3D coordinates of proteins with unknown function is used to annotate these proteins. A very important clue towards function prediction of a protein can be an identification of ligands or small molecules that can bind to the protein. Ligands and other small molecules can be either directly determined within the protein’s 3D structure or a 3D structure of the protein can be used to predict ligand binding sites and thus help to annotate the protein.

A wide range of approaches for protein ligand-binding site prediction have been developed. Table 1 lists those prediction tools that are provided as a web service. Fpocket (2), SiteHound (3), ConCavity (4), POCASA (5), MetaPocket 2.0 (6), FTSite (7) and bSiteFinder (8) support online visualization using Jmol (9), a Java-based molecular structure viewer. However, due to security risks, Java applets are no longer supported in modern web browsers and thus these websites can be considered outdated. A simple solution to the Jmol issue is to use JSmol (10), a JavaScript replacement for Jmol. This is the avenue taken by 3DLigandSite (11), COFACTOR (12, 13), COACH (14) ISMBLAB-LIG (15) and LIBRA (16). Although JSmol supports complex visualization options, it suffers from performance issues due to the inefficiencies introduced when migrating Jmol code from Java to JavaScript. OpenAstex (17), another Java based visualizer, is utilized by Fpocket. However, this project suffers from the same problem as Jmol and seems to be discontinued as we did not manage to find any active resource. Only a few of the web servers support visualization via modern WebGL-based viewers such as LiteMol (18), NGL (19, 20) or PV (21). Specifically, NGL supports visualizations in DoGSite (22) and DeepSite (23). Despite the possibility to view the 3D structures in NGL, DeepSite and DoGSite websites lack the option to customize the protein, ligand and binding sites visualization. Similarly, GalaxySite (24) offers minimal 3D cartoon visualization of the protein and its ligands via the PV viewer. Finally, we have recently developed P2Rank (25), a state-of-the-art method for protein ligand binding site prediction. This work describes PrankWeb, an online web server providing interactive interface for the P2Rank method. The purpose of PrankWeb is to serve as an intuitive tool for ligand binding site prediction and its immediate visual analysis. PrankWeb displays the predictions as a combination of the protein’s 3D structure, sequence and list of binding pockets. It allows users to display protein ligand binding sites and conservation both in the structural and sequence view and customize the style of visualization. PrankWeb’s visualization is based on LiteMol and Protael (26), and thus runs in all modern browsers without additional plugins.

**Table 1.** Availability of existing web-based tools for structure-based ligand binding site prediction introduced since 2009.

| Name | Year | Type | Stand-alone | Online Visualization | Offline Visualization | Source Code |
| --- | --- | --- | --- | --- | --- | --- |
| SiteHound (3) | 2009 | energetic | Yes | Jmol | PyMOL<sup>†</sup>, Chimera<sup>†</sup> | Yes |
| ConCavity (4) | 2009 | conservation | Yes | Jmol | PyMOL | Yes |
| Fpocket (2) | 2010 | geometric | Yes | Jmol, OpenAstex | PyMOL, VMD | Yes |
| 3DLigandSite (11) | 2010 | template | — | JSmol | PyMOL | — |
| POCASA (5) | 2010 | geometric | — | Jmol | — | — |
| DoGSite (22) | 2010 | geometric | — | NGL | — | — |
| MetaPocket 2.0 (6) | 2011 | consensus | — | Jmol | PyMOL | — |
| FTSite (7) | 2012 | energetic | — | Jmol, static | PyMOL | — |
| COFACTOR (12, 13) | 2012, 2017 | template | Yes | JSmol | — | — |
| COACH (14) | 2013 | template | Yes | JSmol | — | — |
| eFindSite (27)* | 2014 | template | Yes | — | PyMOL, VMD, Chimera | Yes |
| GalaxySite (24) | 2014 | template/docking | — | PV, static | — | — |
| bSiteFinder (8) | 2016 | template | — | Jmol | — | — |
| ISMBLab-LIG (15) | 2016 | machine learning | — | JSmol&sequence | — | — |
| LIBRA-WA (16) | 2017 | template | Yes | JSmol | — | — |
| DeepSite (23) | 2017 | machine learning | — | NGL | — | — |
| PrankWeb (P2Rank) | this work | machine learning | Yes | LiteMol & Proteal | PyMOL | Yes |

<sup>†</sup> only data files provided
\*in the process of setting up a new interface

## MATERIALS AND METHODS

### P2Rank

P2Rank (25), the backend of PrankWeb, is a template-free, machine learning-based, ligand binding site prediction method. It employs random forests (28) to predict ligandability of points on the solvent accessible surface of a protein. These points represent potential locations of contact atoms of binding ligands and are described by a feature vector calculated form local geometric neighbourhood. The feature vector consists of physico-chemical and geometrical properties calculated from the surrounding atoms and residues (e.g. hydrophobicity, aromaticity or surface protrusion). PrankWeb also introduces a new model that includes the information derived from sequence evolutionary conservation scores of residues (see Supplementary Information on how the conservation scores are computed). Points with high predicted ligandability are clustered and ranked according to a ranking function based on cumulative score of the cluster.

P2Rank can use different pretrained models with varying feature vectors. PrankWeb exposes two such models: the default P2Rank model (without conservation) and a new model that uses conservation information (P2Rank+Conservation). Both models were trained on relatively small but diverse dataset of protein ligand complexes (25, 29).

As a template-free method, P2Rank does not share the limitation of template based methods which are unable to predict truly novel sites that have no analogues in their template libraries of known protein-ligand complexes. Therefore, P2Rank should be particularly beneficial for predicting novel allosteric sites, for which template based methods would generally be less effective (25). Another advantage of P2Rank is the fact that it is able to work directly with multi-chain structures and predict binding sites formed near the interfaces of chains.

We have compared the predictive performance of the presented tool with several competing algorithms on two datasets: COACH420 (14) containing 420 single-chain complexes and HOLO4K (25) containing 4009 multi-chain structures. Results are shown in Table 2. The default model used by PrankWeb (P2Rank+Conservation) clearly outperforms the other compared methods, and in most cases also the original P2Rank model that does not use conservation. Many of the methods listed in Table 1 are hard to compare on larger datasets as they do not, unlike PrankWeb, expose REST APIs. Thus, batch processing is further hindered by slow running times, delivering results only by email or captcha. Description of evaluation methodology and more detailed results are included in the Supplementary Material.

**Table 2.** Benchmark on COACH420 and HOLO4K datasets.

|  | COACH420 |  | HOLO4K |  |
| --- | --- | --- | --- | --- |
|  | Top-n | Top-(n+2) | Top-n | Top-(n+2) |
| Fpocket 1.0 | 56.4 | 68.9 | 52.4 | 63.1 |
| Fpocket 3.1 | 42.9 | 56.9 | 54.9 | 64.3 |
| SiteHound* | 53.0 | 69.3 | 50.1 | 62.1 |
| MetaPocket 2.0* | 63.4 | 74.6 | 57.9 | 68.6 |
| DeepSite* | 56.4 | 63.4 | 45.6 | 48.2 |
| P2Rank | 72.0 | **78.3** | 68.6 | 74.0 |
| P2Rank+Cons.<sup>†</sup> | **73.2** | 77.9 | **72.1** | **76.7** |

Comparing identification success rate [%] measured by the DCA criterion (distance from pocket center to closest ligand atom) with 4 Å threshold considering only pockets ranked at the top of the list (n is the number of ligands in the considered structure).
\*Failed to produce predictions for some of the input proteins. Here we display success rates calculated only based on subsets of proteins, on which corresponding methods finished successfully.
<sup>†</sup> P2Rank with conservation (the default prediction model of PrankWeb)

Prediction speeds vary heavily among the tools ranging from under 1 second (Fpocket, P2Rank) to more than 10 hours (COACH) for prediction on one protein of average size (2500 atoms). We have previously shown that P2Rank (without conservation) is the second fastest of the available tools (25). PrankWeb does not bring much overhead to prediction speed, however using the model with conservation may take a few minutes if conservation scores need to be calculated from scratch (see Conservation pipeline section in the Supplementary Material).

### Web server

PrankWeb allows users to predict and visualize the predicted protein-ligand binding sites and contrast these with highly conserved areas as well as with actual ligand binding sites.

To carry out the prediction, users can either upload a PDB file or provide a PDB ID in which case PrankWeb will download and store the corresponding PDB file from the PDB database (30). Besides selecting what protein to analyze, users can also specify whether evolutionary conservation should be included in the prediction process, which in turn determines which of the two pretrained models will be used.

Conservation scores are calculated using Jensen-Divergence method (31) from a multiple sequence alignment (MSA) file, which can come from three sources: 1) users can specify their own alignment file, 2) if a protein’s PDB code is provided, PrankWeb uses MSA from the HSSP (32) database, 3) in case no MSA is provided and no MSA is found in HSSP, the MSA is computed using PrankWeb’s own conservation pipeline which utilizes UniProt (33), PSI-Blast (34), MUSCLE (35) and CD-HIT (36). The process is depicted in Figure 2 and described in detail in the Supplementary Material.

After specification of the input, the submitted data is sent via a REST API to the server which starts the prediction pipeline. The user is provided with a URL address where the progress of the prediction process can be tracked and where the results can be inspected once the process finishes.

On the results page, PrankWeb utilizes LiteMol for visualization of the 3D structure information and Protael for sequence visualization. Figure 1 displays predicted binding sites of dasatinib (a drug used for a chromic myelogenous leukemia treatment) bound to the kinase domain of human LCK (PDB ID 3AD5). The sequence and structure plugins are synchronized so that the user can easily locate a sequence position in the structure and vice versa. The sequence view comprises predicted pockets, computed conservation and binding sites (if present in the PDB file). The side panel displays information about identified pockets and a toolbar with the possibility to i) to download all inputs and calculated results, ii) share the results page link or iii) switch between visualization modes. PrankWeb comes with three predefined 3D model renderings: protein surface, cartoon and atoms. Predicted binding sites as well as conservation scores are color coded. If conservation analysis was chosen, the user can contrast the positions of putative active sites with conservation scores of the respective positions. In case the preset modes do not suffice, one can completely customize the 3D visualization using LiteMol’s advanced user interface or using the PyMOL visualization script for offline inspection.

**Figure 1.**
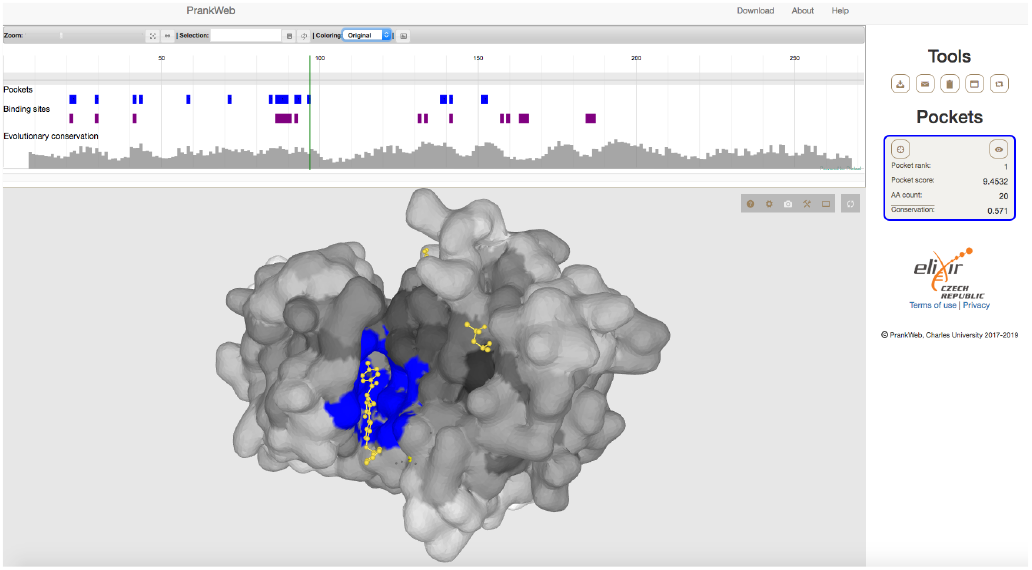
An example of PrankWeb output. The figure shows a prediction of ligand binding sites (blue colour) on a surface of human Lck kinase (3AD5). The actual ligand binding pose of dasatinib is shown in yellow. The second small molecule in the figure is dimethyl sulfoxide. The figure also shows a sequence view on the protein with indicated binding sites and conservation scores (top panel). The right panel shows a summary of the binding sites and provides tools to modify the view or to download the results.

**Figure 2.**
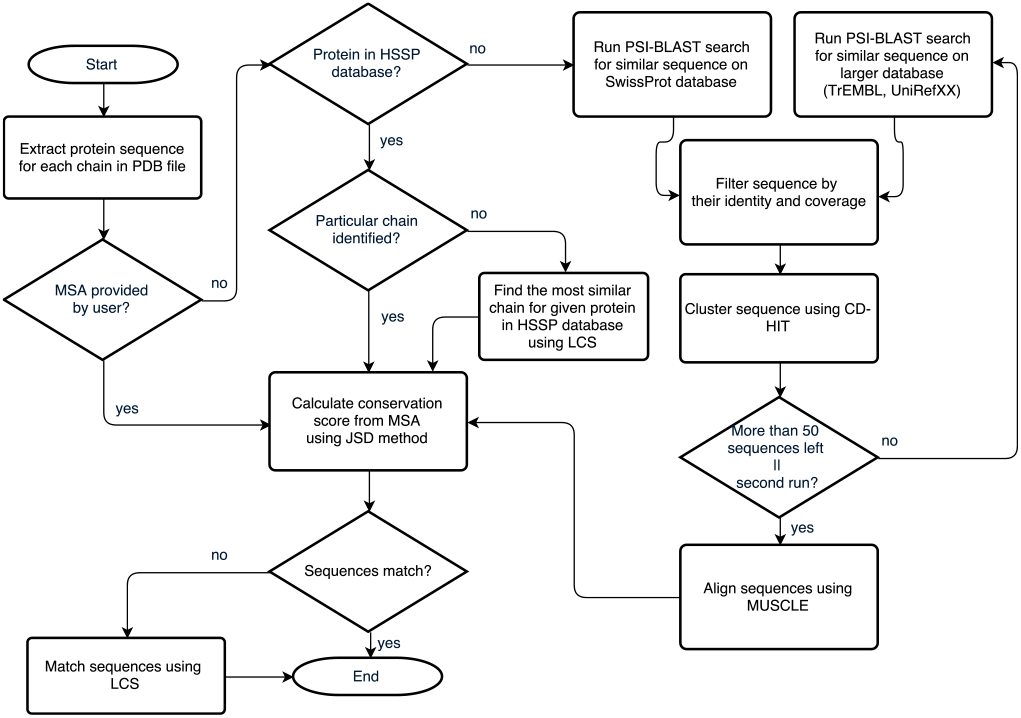
Diagram of conservation loading workflow and conservation pipeline.

PrankWeb consists of a Java backend, REST API and a Typescript frontend. The backend is based on WildFly (37) web server and P2Rank application. The frontend utilizes Protael, LiteMol and Bootstrap.js libraries to provide interactive user interface on top of the REST API. All source code is available under the Apache License 2.0 at GitHub (https://github.com/jendelel/PrankWebApp). The GitHub website also includes documentation for other developers on how to use our REST API as well as how to deploy their own instance of the server.

## DISCUSSION

PrankWeb has proven to provide correct predictions even in cases where other methods have failed. Nghan et al. (7) mentioned three cases where their method FTSite was unable to identify the ligand binding site with their best ranked prediction. PrankWeb correctly identified the binding site as the best ranked both in apo and holo structure in all three cases. Namely, in the case of glucose/galactose receptor (1GCG, 1GCA), purine nucleoside phosphorylase (1ULA,1ULB) and finally also in the case of mouse FV fragment of antibody (1A6U,1A6W). Figure 3 shows the predicted ligand binding site of holo structure (1GCA) on the interface of two imunoglobulin subunits together with experimentaly solved structure of 4-HYDROXY-5-IODO-3-NITROPHENYLACETYL-EPSILON-AMINOCAPROIC ACID ANION (NIP). The 3D structure of NIP appears in the PDB just once, which makes it difficult to train its binding.

**Figure 3.**
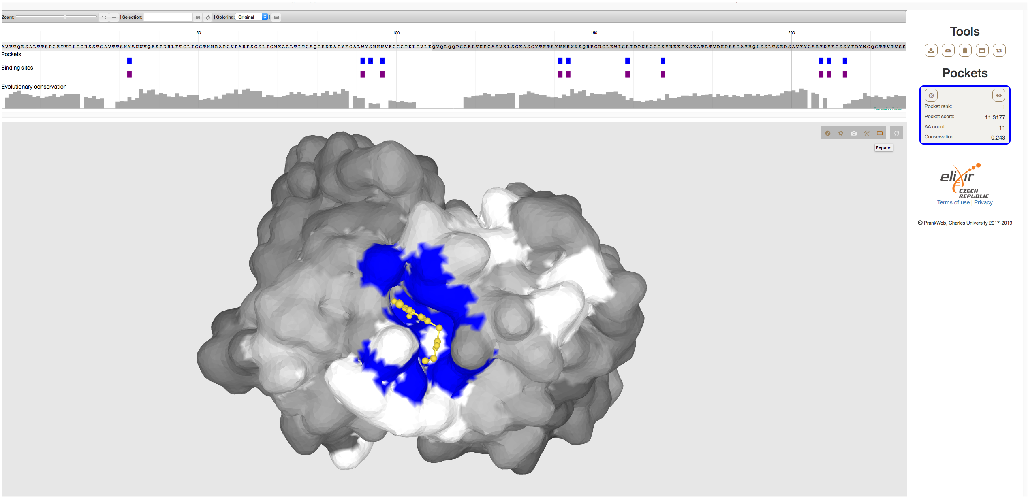
Prediction a difficult pocket. Authors of FTSite method described three structures, where their method failed. This figure shows a PrankWeb prediction for one of them - a structure of mouse immunoglobulin (1a6w). The prediction is indicated by a blue colour and the actual ligand is in yellow.

We should highlight that none of the models employed by PrankWeb was trained on metallic ion ligands and would not in general be good at finding their binding sites. Such task would be better served by a model trained on a specialized dataset (which is possible to do with P2Rank).

## CONCLUSION

We presented PrankWeb, a new web interface for the state-of-the-art ligand binding prediction method P2Rank. PrankWeb allows users to quickly carry out a prediction and visually inspect the results. PrankWeb also contains a pipeline for computation of conservation scores, which are included in the ligand binding site prediction as well as the structure-sequence visualization of the results. While PrankWeb provides a user-friendly interface, it also serves a REST API, enabling developers to use PrankWeb as a service. Both PrankWeb and P2Rank are open sourced on GitHub and freely available.

## ACKNOWLEDGEMENTS

Access to computing and storage facilities owned by parties and projects contributing to the National Grid Infrastructure MetaCentrum provided under the programme “Projects of Large Research, Development, and Innovations Infrastructures” (CESNET LM2015042), is greatly appreciated.

## FUNDING

This work was supported by ELIXIR CZ research infrastructure project (MEYS Grant No: LM2015047) and by the Grant Agency of Charles University (project Nr. 1556217).

## Conflict of interest statement

None declared.

