## Supplementary material for "PrankWeb: web server for ligand binding-site prediction and visualization"

### CONSERVATION PIPELINE

Conservation scores for PrankWeb are computed from multiple sequence alignment (MSA). MSA for a particular sequence can be acquired from an HSSP database (1) or calculated from a set of sequences using bioinformatics tools. Moreover, PrankWeb also allows users to upload their own MSA for each chain.

If the HSSP database contains the protein of interest and no chain for the particular ID was found, PrankWeb takes the chain with the longest common subsequence. In case the protein is not present in HSSP and the user did not provide the MSA for that protein, homology pipeline is invoked to obtain an MSA. The main idea of the pipeline (inspired by ConSurfDB (2)) is based on querying databases for similar sequences to the input sequence. The decision making process for calculating conservation scores is illustrated in Figure 2 of the main article. It takes a protein sequence in FASTA format as input and outputs a tab-separated file with conservation scores, which is the result of the Jensen-Shannon divergence method for calculating the conservation scores from multiple sequence alignment. (3)

The pipeline proceeds as follows:

1. SwissProt is queried for similar protein sequences using PSI-BLAST (4) with e-value=$10^{-5}$. ConSurfDB uses the same e-value.
2. The sequences that are too similar or too different than our query sequence are filtered out.
3. Then CD-HIT (5) is run with default parameters to cluster the sequences and outputs a non-redundant representative sequence list.
4. If less than 50 sequences are left, we repeat the steps 1–3 on, the larger database, UniRef90 (6).
5. Sequences are aligned using MUSCLE (7).
6. At this point, we have a multiple sequence alignment and can calculate the conservation score using the Jensen-Shannon divergence method (3).

### EVALUATION METHODOLOGY

To evaluate predictive performance of PrankWeb we have used the same methodology that was used in original P2Rank article (8). It is based on ligand-centric counting and the DCA (distance between the center of the pocket and any ligand atom) pocket identification criterion with 4 Å threshold. Ground-truth binding sites are defined by ligands present in evaluation datasets. Every structure in a dataset can contain more than one relevant ligand (see below) and for every relevant ligand, its binding site must be correctly predicted for a method to achieve 100% identification success rate on the given dataset. Every relevant ligand contributes with equal weight toward the final success rate. The output of prediction methods is a ranked list of several putative binding sites, but during evaluation only those ranked at the top are considered. We use Top-$n$ and Top-($n$+2) rank cutoffs where for every evaluated protein structure $n$ is the number of relevant ligands in this structure (i.e. for proteins that have only one ligand this corresponds to the usual Top-1 and Top-3 cutoffs and for proteins with 2 ligands to Top-2 and Top-4 cutoffs). This evaluation methodology is the same as the one that was used in the only independent benchmark of ligand binding site prediction algorithms to date (9).

#### Relevant Ligands

P2Rank is focused on predicting binding sites for biologically relevant ligands and PDB files in considered datasets often contain ligands (i.e. HET groups) that are not relevant. To determine which ligands in benchmark datasets are relevant we use a custom filter and alternatively the binding MOAD (10) database.

In addition to biologically relevant ligands, PDB files contain a variety of other HET groups like solvents, salt and misplaced groups (that are not in contact with the protein). Instead of declaring only one ligand as relevant for every file in a dataset (as was done in other ligand binding site prediction studies), we determine relevant ligands by a filter. Ligands that are considered relevant must comply to these conditions:

- Number of ligand atoms is greater or equal than 5.

† The work has been done while studying at Charles University.

**Table 1.** Benchmark on COACH420, COACH420(Mlig), HOLO4K and HOLO4K(Mlig) datasets.

| | COACH420 | | COACH420(Mlig) | | HOLO4K | | HOLO4K(Mlig) | |
|---|---|---|---|---|---|---|---|---|
| | Top-n | Top-(n+2) | Top-n | Top-(n+2) | Top-n | Top-(n+2) | Top-n | Top-(n+2) |
| Fpocket 1.0 | 56.4 | 68.9 | 57.4 | 70.4 | 52.4 | 63.1 | 56.9 | 70.3 |
| Fpocket 3.1 | 42.9 | 56.9 | 43.1 | 56.3 | 54.9 | 64.3 | 57.4 | 69.1 |
| SiteHound* | 53.0 | 69.3 | 51.0 | 67.7 | 50.1 | 62.1 | 53.1 | 67.8 |
| MetaPocket 2.0* | 63.4 | 74.6 | 62.2 | 73.3 | 57.9 | 68.6 | 62.3 | 75.2 |
| DeepSite* | 56.4 | 63.4 | 54.5 | 61.6 | 45.6 | 48.2 | 50.8 | 54.4 |
| P2Rank | 72.0 | **78.3** | **71.2** | **76.5** | 68.6 | 74.0 | 73.7 | 80.9 |
| P2Rank+Conservation† | **73.2** | 77.9 | 70.9 | 75.1 | **72.1** | **76.7** | **77.2** | **83.3** |

Comparing identification success rate [%] measured by the DCA criterion (distance from pocket center to closest ligand atom) with 4 Å threshold considering only pockets ranked at the top of the list (n is the number of ligands in the considered structure).

*Failed to produce predictions for some of the input proteins. Here we display success rates calculated only based on subsets of proteins, on which corresponding methods finished successfully. Detailed, pairwise comparison with P2Rank on the exact subsets can be found in the Supplementary Information of P2Rank article (8).

† P2Rank with conservation (the default prediction model of PrankWeb)

- Distance from any atom of the ligand to the closest protein atom is at least 4 Å (to remove "floating" HET groups present in some structures).
- Distance form the center of the mass of the ligand to the closest protein atom is not greater than 5.5 Å (to remove ligands that "stick out").
- Name of the PDB group is not on the list of ignored groups: `(HOH, DOD, WAT, NAG, MAN, UNK, GLC, ABA, MPD, GOL, SO4, PO4)`.

Choosing relevant ligands in this particular way is admittedly arbitrary. In order to make sure our results are robust with respect to the exact way relevant ligands are determined, we have created a versions of COACH420 and HOLO4K datasets where relevant ligands are determined in a different way. Binding MOAD (10) release 2013, a database of biologically relevant ligands in PDB, was used to determine relevant ligands in resulting datasets COACH420(Mlig) and HOLO4K(Mlig). PDB files that have no entry in MOAD were removed from the new datasets.

It should be noted that the notion of a biologically relevant ligand does not have a widely accepted definition. There are other databases that purportedly collect only biologically relevant ligand interactions from the PDB (e.g. BioLiP (11), PDBbind (12)) that use different criteria for accepting particular ligand as biologically relevant (with MOAD being the strictest of them, for example, by not accepting any small ions). For a discussion on the caveats of determining biologically relevant ligands see (11).

### Datasets

All datasets used to train and optimize our models and produce presented results are available on GitHub http://github.com/rdk/p2rank-datasets and described in detail in P2Rank paper (8).

P2Rank was trained on the CHEN11 dataset (both models employed by PrankWeb: with and without conservation) and various parameters of the algorithm were optimized with respect to the results on the JOINED dataset (8), that was used as a development/validation dataset. For future benchmarks we note that results on proteins from those datasets would not represent an unbiased estimate of P2Rank's performance.

### ADDITIONAL RESULTS

Table 1 is an extended version of the results table from the main article which includes results on *(Mlig) versions of datasets where relevant ligands were determined differently (see Relevant Ligands section). It shows that our results are robust with respect to the particular way relevant ligands are determined. New P2Rank model with conservation seems to perform slightly worse on COACH420 dataset but substantially better on larger HOLO4K dataset. Table 2 shows average numbers of predicted sites for each method. P2Rank+Conservation in general predicts fewer but more relevant sites than the original P2Rank model.

The results were taken from (8) and we performed new benchmark experiments for Fpocket 3.1 and P2Rank+Conservation. Results of Fpocket 3.1 correspond to the 3.1.2 version downloaded and compiled from GitHub (https://github.com/Discngine/fpocket), run with default parameters.

**Table 2.** Number of predicted binding sites and dataset statistics.

| | COACH420 | HOLO4K |
|---|---|---|
| Proteins | 420 | 4009 |
| Avg. protein atoms | 2179 | 3908 |
| Avg. ligands | 1.2 | 2.4 |
| Fpocket 1.0 | 14.6 | 27.0 |
| Fpocket 3.1 | 13.9 | 16.0 |
| SiteHound | 66.2 | 99.5 |
| MetaPocket 2.0 | 6.3 | 6.4 |
| DeepSite | 3.2 | 2.8 |
| P2Rank | 6.3 | 12.6 |
| P2Rank+Conservation | 3.4 | 7.7 |

Displayed is the average total number of binding sites predicted per protein by each method on a given dataset.
